# Boltz-1 Democratizing Biomolecular Interaction Modeling

**DOI:** 10.1101/2024.11.19.624167

**Authors:** Jeremy Wohlwend, Gabriele Corso, Saro Passaro, Noah Getz, Mateo Reveiz, Ken Leidal, Wojtek Swiderski, Liam Atkinson, Tally Portnoi, Itamar Chinn, Jacob Silterra, Tommi Jaakkola, Regina Barzilay

**Affiliations:** MIT CSAIL; MIT Jameel Clinic; Genesis Research, a part of Genesis Therapeutics; CHARM Therapeutics

## Abstract

Understanding biomolecular interactions is fundamental to advancing fields like drug discovery and protein design. In this paper, we introduce Boltz-1, an open-source deep learning model incorporating innovations in model architecture, speed optimization, and data processing achieving AlphaFold3-level accuracy in predicting the 3D structures of biomolecular complexes. Boltz-1 demonstrates a performance on-par with state-of-the-art commercial models on a range of diverse benchmarks, setting a new benchmark for commercially accessible tools in structural biology. Further, we push the boundary of capabilities of these models with Boltz-steering, a new inference time steering technique that is able to fix hallucinations and non-physical predictions from the models. By releasing the training and inference code, model weights, datasets, and benchmarks under the MIT open license, we aim to foster global collaboration, accelerate discoveries, and provide a robust platform for advancing biomolecular modeling.

## 1 Overview

Biomolecular interactions drive almost all biological mechanisms, and our ability to understand these interactions guides the development of new therapeutics and the discovery of disease drivers. In 2020, AlphaFold2 [Jumper et al., 2021] demonstrated that deep learning models can reach experimental accuracy for single-chain protein structure prediction on a large class of protein sequences. However, a critical question about modeling biomolecular complexes in 3D space remained open.

In the past few years, the research community has made significant progress toward solving this pivotal problem. In particular, the use of deep generative models has proven to be effective in modeling the interaction between different biomolecules with diffDock [Corso et al., 2022] showing significant improvements over traditional molecular docking approaches and, most recently, alphaFold3 [Abramson et al., 2024] reaching unprecedented accuracy in the prediction of arbitrary biomolecular complexes.

In this manuscript, we present Boltz-1, the first fully commercially accessible open-source model reaching alphaFold3 reported levels of accuracy. By making the training and inference code, model weights, datasets, and benchmarks freely available under the MIT license, we aim to empower researchers, developers, and organizations around the world to experiment, validate, and innovate with Boltz-1. At a high level, Boltz-1 follows the general framework and architecture presented by Abramson et al. [2024], but it also presents several innovations which include:

1. New algorithms to more efficiently and robustly pair MSAs, crop structure at training time, and condition predictions on user-defined binding pockets;
2. Changes to the flow of the representations in the architecture and the diffusion training and inference procedures;
3. Revision of the confidence model both in terms of architectural components as well as the framing of the task as a fine-tuning of the model”s trunk layers.

In the following sections, we detail these changes as well as benchmark the performance of Boltz-1 with other publicly available models. Our experimental results show that Boltz-1 delivers performance on par with the state-of-the-art commercial models on a wide range of structures and metrics.

Further, we tackle one of the biggest outstanding challenges with machine learning based structure prediction methods, their frequent lack of respect for physical laws observed, for example, in mistakes in properties like internal geometries, chirality, and steric clashes as well as hallucinations like overlapping chains. We propose a new inference time technique, Boltz-steering, which is able to solve almost all these physical issues while maintaining the model accuracy. We refer to the Boltz-1 model with the steering as Boltz-1x.

Given the dynamic nature of this open-source project, this manuscript and its linked GitHub repository^1^ will be regularly updated with improvements from our core team and the community. We aspire for this project and its associated codebase to serve as a catalyst for advancing our understanding of biomolecular interactions and a driver for the design of novel biomolecules.

## 2 Data pipeline

Boltz-1 operates on proteins represented by their amino acid sequence, ligands represented by their smiles strings (and covalent bonds), and nucleic acids represented by their genomic sequence. This input is then augmented by adding multiple sequence alignment (MSA) and predicted molecular conformations. Unlike AlphaFold3, we do not include input templates, due to their limited impact on the performance of large models.

In this section, we first outline how the structural training data, as well as the MSA and conformer, were obtained and describe the curation of our validation and test sets. Then, we describe three important algorithmic developments applied to data curation and augmentation that we find to be critical:

1. A new algorithm to pair MSAs for multimeric protein complexes from taxonomy information (2.3).
2. A unified cropping algorithm that combines the spatial and contiguous cropping strategies used in previous work (2.4).
3. A robust pocket-conditioning algorithm tailored to common use cases (2.5).

### 2.1 Data source and processing

#### PDB structural data

For training we use all PDB structures [Berman et al., 2000] released before 2021-09-30 (same training cut-off date as AlphaFold3) and with a resolution of at least 9Å. We parse the Biological Assembly 1 from these structures from their mmCIF file. For each polymer chain, we use the reference sequence and align it to the residues available in the structure. For ligands, we use the CCD dictionary to create the conformers and to match atoms from the structure. We remove leaving atoms when (1) the ligand is covalently bound and (2) that atom does not appear in the PDB structure. Finally, we follow the same process as AlphaFold3 for data cleaning, which includes the ligand exclusion list, the minimum number of resolved residues, and the removal of clashing chains.

#### MSA and molecular conformers

We construct MSAs for the full PDB data using the `colabfold_search` tool [Mirdita et al., 2022] (which leverages MMseqs2 [Steinegger and Söding, 2017]), using default parameters (versions: `uniref30_2302, colabfold_envdb_202108`). We then assign taxonomy labels to all UniRef sequences using the taxonomy annotation provided by UniProt [Consortium, 2015]. For the initial molecular conformers that are provided to the model, we pre-compute a single conformer for all CCD codes using the RDKit’s ETKDGv3 [Wang et al., 2022].

#### Structure prediction training pipeline

We train the structure prediction model (see Section 3.2 for details of the confidence model training) for a total of 68k steps with a batch size of 128. During the first 53k iterations, we use a crop size of 384 tokens and 3456 atoms and draw structures equally from the PDB dataset and the OpenFold distillation dataset (approximately 270K structures, using the MSAs they provided) [Ahdritz et al., 2024]. For the last 15k iterations, we only sampled from the PDB structures and had a crop size of 512 tokens and 4608 atoms. As a comparison AlphaFol3 trained a similar architecture for nearly 150k steps with a batch size of 256, which required approximately four times the computing time. We attribute some of this drastic reduction to the various innovations we detail in the remainder of this section and the next.

### 2.2 Validation and test sets curation

To address the absence of a standardized benchmark for all-atom structures, we are releasing a new PDB split designed to help the community converge on reliable and consistent benchmarks for all-atom structure prediction tasks. Our training, validation and test splitting strategy largely follows Abramson et al. [2024]. We first cluster the protein sequences in PDB by sequence identity with the command `mmseqs easy-cluster … --min-seq-id 0.4` interaction between different biomolecules with [Hauser et al., 2016]. Then, we select all structures in PDB satisfying the following filters:

1. Initial release date is before 2021-09-30 (exclusive) and 2023-01-13 (inclusive).
2. Resolution is below 4.5 Å.
3. All the protein sequences of the chains are not present in any training set clusters (i.e. before 2021-09-30).
4. Either no small-molecule is present, or at least one of the small-molecules exhibits a Tanimoto similarity of 0.8 or less to any small-molecule in the training set. Here, a small-molecule is defined as any non-polymer entity containing more than one heavy atom and not included in the ligand exclusion list.

This yields 1728 structures, which we further refine through the following steps:

1. Retaining all the structures containing RNA or DNA entities. (126 structures)
2. Iteratively adding structures containing small-molecules or ions under the condition that all their protein chains belong to new unseen clusters (330 additional structures)
3. Iteratively adding multimeric structures under the condition that all the protein chains belong to new unseen clusters. These are further filtered by randomly keeping only 50% of the passing structures. (231 additional structures)
4. Iteratively adding monomers under the condition that their chain belongs to a new unseen cluster. These are further randomly filtered out by keeping only 30% of the passing structures. (57 additional structures)

This results in a total of 744 structures. Finally, we retain the structures with at most 1024 residues in the valid protein/RNA/DNA chains, finishing with a total of 553 validation set structures.

The test set is created using the same procedure described above with the following differences: for protein and ligand similarity exclusion we consider all structures released before 2023-01-13 (which include all training and validation sets), we filter to structures released after 2023-01-13 and the final size filter to structures between 100 and 2000 total residues. The resulting final test set size is 593.

### 2.3 Dense MSA pairing algorithm

Multiple sequence alignments uncover amino acids that co-evolved throughout evolution, and therefore are likely close to each other in physical space. However, extracting such signals for protein-protein interactions poses a greater challenge, as most proteins are sequenced or reported individually. To approximate these pairings, researchers have leveraged the taxonomy information frequently associated with sequences. In Algorithm 3, we present a method for pairing MSAs using taxonomy in a manner that preserves MSA density (a critical factor, as model complexity scales linearly with the number of MSA rows) while balancing the trade-off between the signal derived from paired sequences and the sequence redundancy within each chain.

### 2.4 Unified cropping algorithm

In order to efficiently train on complexes with variable size, methods like alphaFold2 and AlphaFold3 crop the structures during training to a fixed maximum number of atoms, residues, or tokens. The most common techniques to perform such crops are (1) contiguous, where tokens are chosen to be consecutive residues in a biomolecular sequence (or entire molecules), and (2) spatial crops, where tokens are chosen purely depending on their distance from a center token. Each of these two has its advantages and provides different training signals to the models, therefore they are often used in combination as done, for example, by Abramson et al. [2024].

We argue, however, that these are two extremes and it is useful to train the model on a more diverse range of cropping strategies. To this end, we define a new cropping algorithm which directly interpolates between spatial and contiguous strategies. The algorithm, formalized in Algorithm 4, revolves around the definition of neighborhoods, that characterize contiguous portions of sequences of a particular length (or entire non-polymer entities) around a specific token. Neighborhoods are incrementally added to the crop depending on the distance of their central token from the chosen center of the crop. If the size of the neighborhoods is chosen to be zero, this strategy translates into spatial cropping, whereas if the size is half of the maximum token budget, this strategy translates into continuous cropping. In our experiments, we find it beneficial to randomly sample the neighborhood size uniformly between zero and 40 tokens for every training sample.

### 2.5 Robust pocket-conditioning

In many real-world scenarios, researchers have prior knowledge of the protein”s binding pocket. Therefore, it is valuable to enable the model to condition on the pocket information. AlphaFold3 explored pocket-conditioned generation by fine-tuning the model to include an additional token feature for all the pocket-ligand pairs, where the pocket is defined as any residue with heavy atoms within 6Å of the ligand. While effective, this design has some limitations. It requires maintaining two models, one with and one without pocket conditioning, and it assumes the specification of all residues within 6Å. This assumption may not align with realistic scenarios, where users might only know key residues, and the full set of interacting residues is highly dependent on the ligand pose, which is often unknown.

To address these challenges, we implement a different strategy for pocket conditioning, designed to (1) retain a single unified model, (2) ensure robustness to a partial specification of interacting residues, and (3) enable interaction site specification for polymer binders such as proteins or nucleic acids. During training, we incorporate pocket information for a randomly selected binder in 30% of iterations. For these cases, we draw the (maximum) number of pocket residues to reveal from a geometric distribution and randomly select residues from those with at least one heavy atom within 6Å of the binder. This information is then encoded as an additional one-hot token feature provided to the model. The training process for this pocket-conditioning approach is described in detail in Algorithm 5.

## 3 Modeling

For the model architecture and training, we started by reproducing AlphaFold3 as described in the supplementary material of Abramson et al. [2024]. AlphaFold3 is a diffusion model that uses a multi-resolution transformer-based model for the denoising of atom coordinates. The model operates at two levels of resolution: heavy atoms and tokens. Tokens are defined as amino acids for protein chains, nucleic acid bases for RNA and DNA, and individual heavy atoms for other molecules and modified residues or bases.

On top of the denoising transformer, critically, AlphaFold3 also employs a central trunk architecture that is used to initialize tokens” representations and determine the denoising transformer”s attention pair bias. This trunk is computationally expensive due to its use of token pairs as fundamental “computational token” and its axial attention operations on these pair representations which results in a complexity that scales cubically with the number of input tokens. To make such encoding computationally tractable, the trunk is set to be independent of the specific diffusion time or input structure such that it can be run only once per complex.

Starting from this architecture, we designed and tested a number of potential alternative approaches. In the following sections, we describe the ones that yielded improvements and were therefore adopted into Boltz-1.^2^ Because of the significant computational budget required to train a full-sized model, we tested these changes on a smaller-sized architecture at different points of our development process. We expect our observations to hold for the final full-size model, but cannot present direct ablation studies.

### 3.1 Architectural modifications

#### MSA module

We find it beneficial to reorder the operations performed in the `MSAModule` (AlphaFold3 Algorithm 8) to better allow the updates on the single and pair representations to feed to one another. In particular, we change the order^3^ of its operations from:

~~~
OuterProductMean, PairWeightedAveraging, MSATransition, TriangleUpdates, PairTransition
~~~

to:

~~~
PairWeightedAveraging, MSATransition, OuterProductMean, TriangleUpdates, PairTransition.
~~~

Note that `OuterProductMean` propagates information from the single to the pair representation, so we now allow the single representations learned in the `MSATransition` to directly propagate to the pair representation.

#### Transformer layer

Abramson et al. [2024] presents an unusual order of operations in their `DiffusionTransformer` layers where hidden representations are updated as (alphaFold3 Algorithm 23):

~~~
a ← AttentionPairBias(a) + ConditionedTransitionBlock(a).
~~~

This has two issues (1) it lacks residual connections that may make backpropagation more complex and (2) it does not allow for the transformation learned in the `AttentionPairBias` to be fed in the `ConditionedTransitionBlock` at the same block. We found it to be beneficial to apply the following transformation order:

~~~
a ← a + AttentionPairBias(a)
a ← a + ConditionedTransitionBlock(a).
~~~

### 3.2 Training and inference procedures

#### Kabsch diffusion interpolation

A key change between AlphaFold2 and AlphaFold3 was the non-equivariance of the denoising model of AlphaFold3 (compared to the equivariant IPA-based structure module of AlphaFold2) to rotations and translations. To encourage the robustness of the denoising model to such transformations their input is randomly translated and rotated before the denoising at training and inference times. To further reduce the variance of the denoising loss with respect to these variations, Abramson et al. [2024] use a rigid alignment between the predicted denoising coordinates and the true coordinates before computing the MSE loss.

However, we argue that on its own this procedure is theoretically problematic. One can define simple functions that would achieve zero rigid aligned MSE loss during training, but completely fail to sample realistic poses at inference time. For example, consider a model trying to fit a given structure with coordinates *x*. Let”s assume that for any noised structure within some reasonable noising perturbation (e.g. Δ = 10*σ*_*t*_), the model always predicts *x*:

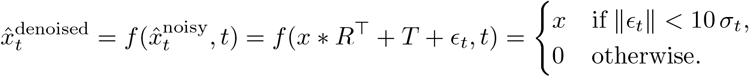

where *R* and *T* are respectively a random rotation matrix and a random translation vector, *ϵ*_*t*_ and *σ*_*t*_ represent respectively the random noise and noise standard deviation for some diffusion time *t*. This model will have a loss approaching zero during training (one will never sample something beyond 10 standard deviations, and one could make this Δ arbitrarily large). However, when used at inference time, this model will consistently go out of distribution (and therefore predict a zero vector). This is because at low noise levels the interpolation between the current randomly rotated 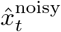 and the predicted 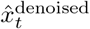 may lead to a pose 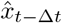 that is very far from *x* and will fall beyond the 10 *σ*_*t*_ mark. Figure 2 shows a graphical representation of this issue.

**Figure 1:**
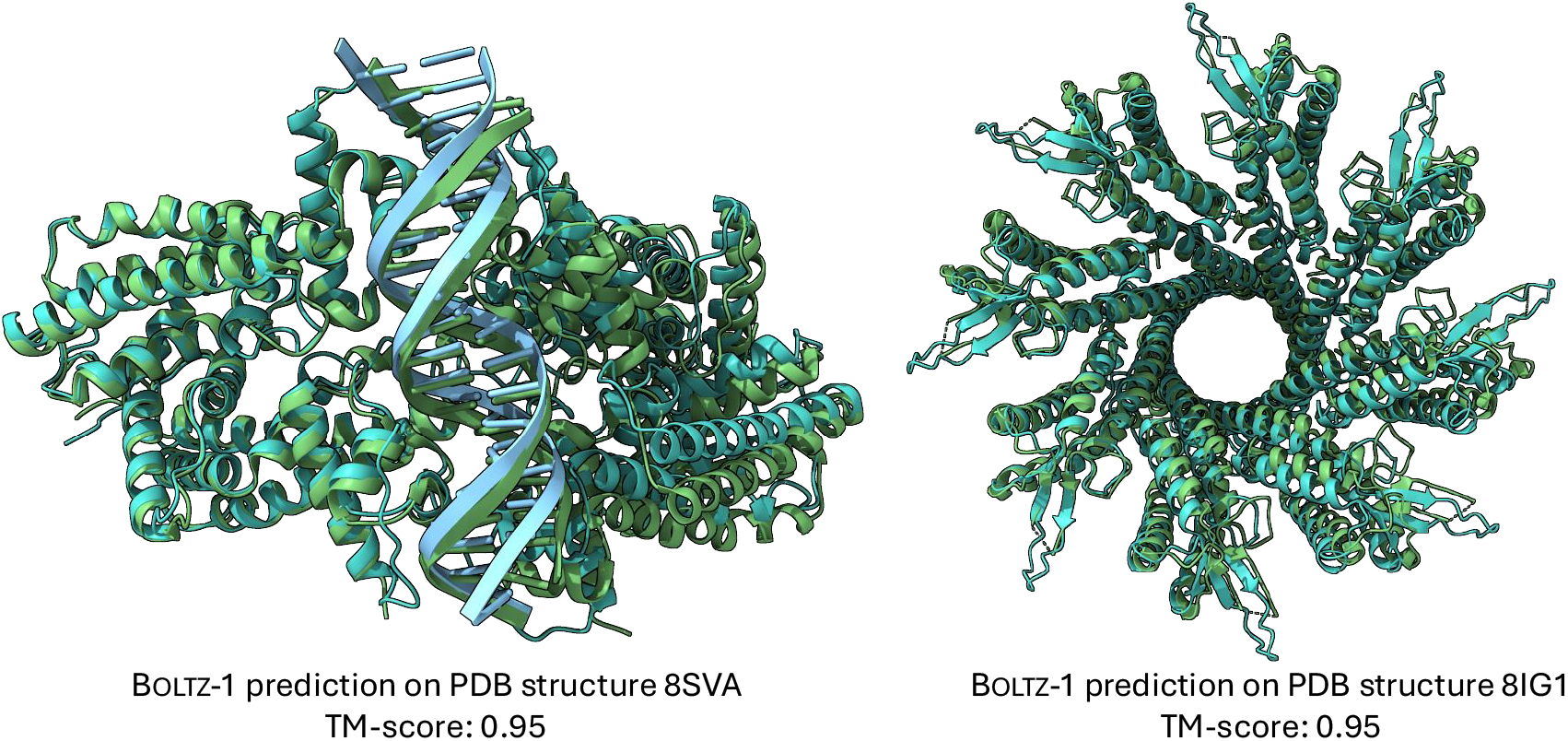
Example predictions of Boltz-1 on targets from the test set.

**Figure 2:**
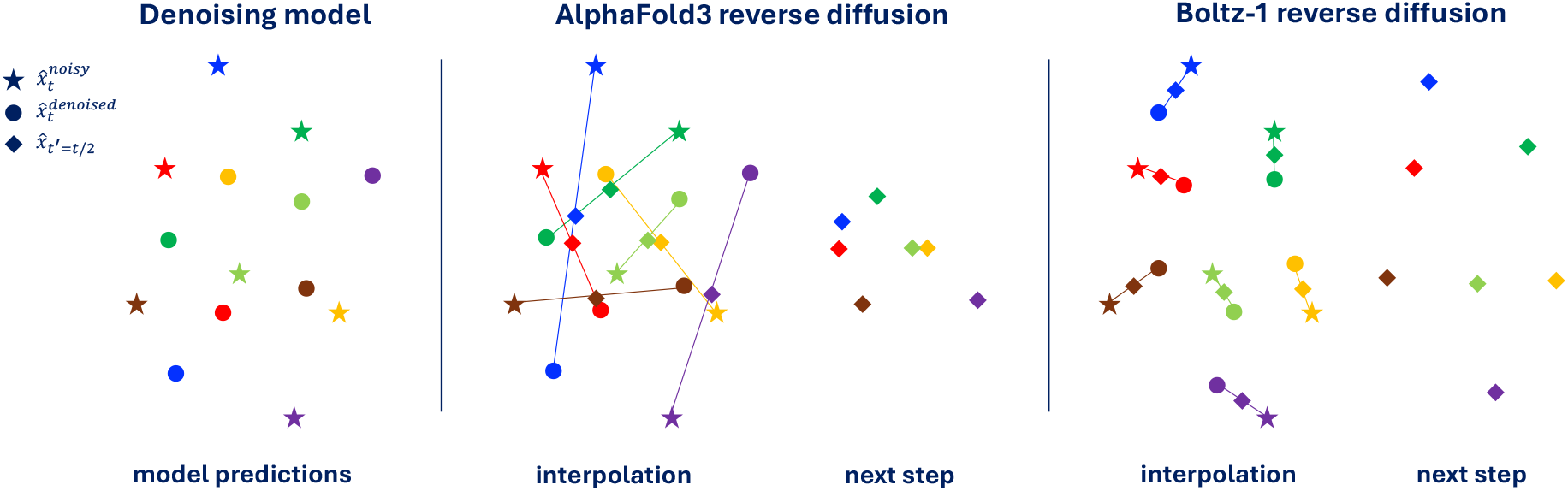
2D representation of the difference between AlphaFold3 reverse diffusion and Boltz-1 reverse diffusion with our Kapsch interpolation. Colors indicate correspondence between different points. Even though the prediction of the denoising model is “perfect” according to the aligned MSE loss, the unaligned interpolation may lead to poor structures fed to the next reverse diffusion step.

We overcome this issue by adding a rigid alignment with Kabsch algorithm after every step during the inference procedure before 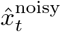 and 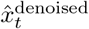 are interpolated (see Figure 2 for a visual explanation). Informally, our diffusion interpolation operates on the minimal projection between noisy and denoised structures, guaranteeing, under the assumption of a Dirac distribution, that the interpolated structure is more similar to the denoised sample than the noisy structure. Empirically, we note that this change to the reverse diffusion has a bigger effect when training models on subsets of the full data where the model is more likely to overfit, on the other hand, the final Boltz-1 seems to largely denoising close to the projection making the Kapsch alignment not critical.

#### Diffusion loss weighting

For the weighting of the diffusion loss we use 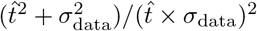 in line with the EDM framework [Karras et al., 2022], rather than 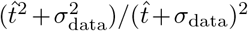 (AlphaFold3 Section 3.7.1 Eq. 6).

### 3.3 Confidence model

AlphaFold3 trains the confidence model alongside the trunk and denoising models while, however, cutting all the gradients going from the confidence task to the rest of the model. Instead, training structure prediction and confidence models separately allowed us to disentangle experiments on each component and make several important improvements to the confidence prediction task.

In AlphaFold3 the architecture of the confidence model is composed of four PairFormer layers that take as input the final single and pair token representations from the model trunk as well as an encoding of the token pairwise distances predicted by the reverse diffusion. These four layers are followed by linear projections trained to predict whether each atom is resolved in the crystal structure, per-atom LDDT and per-token pair PAE and PDE.

#### Trunk architecture and initialization

We noticed that, at a high level, the input-output composition of the confidence model is similar to that of the trunk. The trunk also takes as input its own final representations (through recycling) and outputs expressive representations used by the denoising model. Therefore, inspired by the way that researchers in the large language model community have been training reward models by fine-tuning the “trunk” of their pretrained generative models [Touvron et al., 2023], we define the architecture of our confidence model to contain all the components of the trunk and initialize its representation to the trained trunk weights. Hence, our confidence model presents an `AtomAttentionEncoder`, an `MSAModule`, and a `PairFormerModule` with 48 layers. In addition, we still integrate the predicted conformation as an encoding of the pairwise token distance matrix and decode the confidence with linear layers on the final PairFormer representation.

#### Diffusion model features

We feed to the confidence model not only the representations coming from the trunk but also a learned aggregation of the final token representation at each reverse diffusion step. These representations are aggregated through the reverse diffusion trajectory with a time-conditioned recurrent block and then fed concatenated to the trunk token-level features at the start of the confidence model. We further modify the way that token-level features are fed to the pairwise representations adding an element-wise multiplication of linearly transformed token-level features.

#### Overall procedure and training

We detail our new full inference procedure in Algorithm 1 and provide a schematic representation in Figure 3. To train the confidence model we initialize all the components borrowed by the trunk to the final trunk weights (from the exponentially moving average) and initialize the weights of all the other components of the network randomly but with zeroed final layers not to perturb initial rich representation from the pretrained weights.

**Figure 3:**
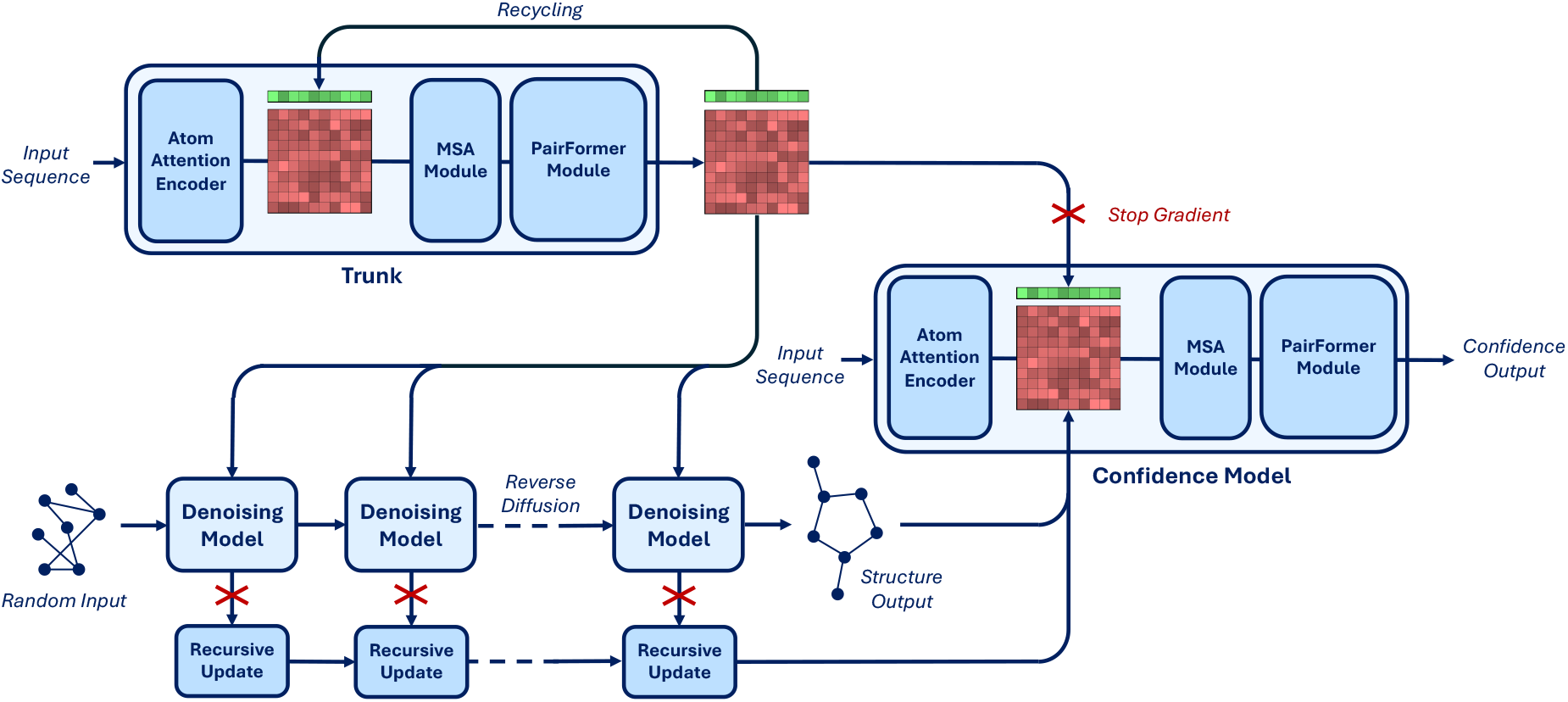
Diagram of the architecture of Boltz-1. The critical difference with AlphaFold3 lies in the confidence model, which now not only has a `PairFormerModule` but follows a full trunk composition and is fed features coming from the denoising model through the recursive updates.

##### Algorithm 1: Confidence model

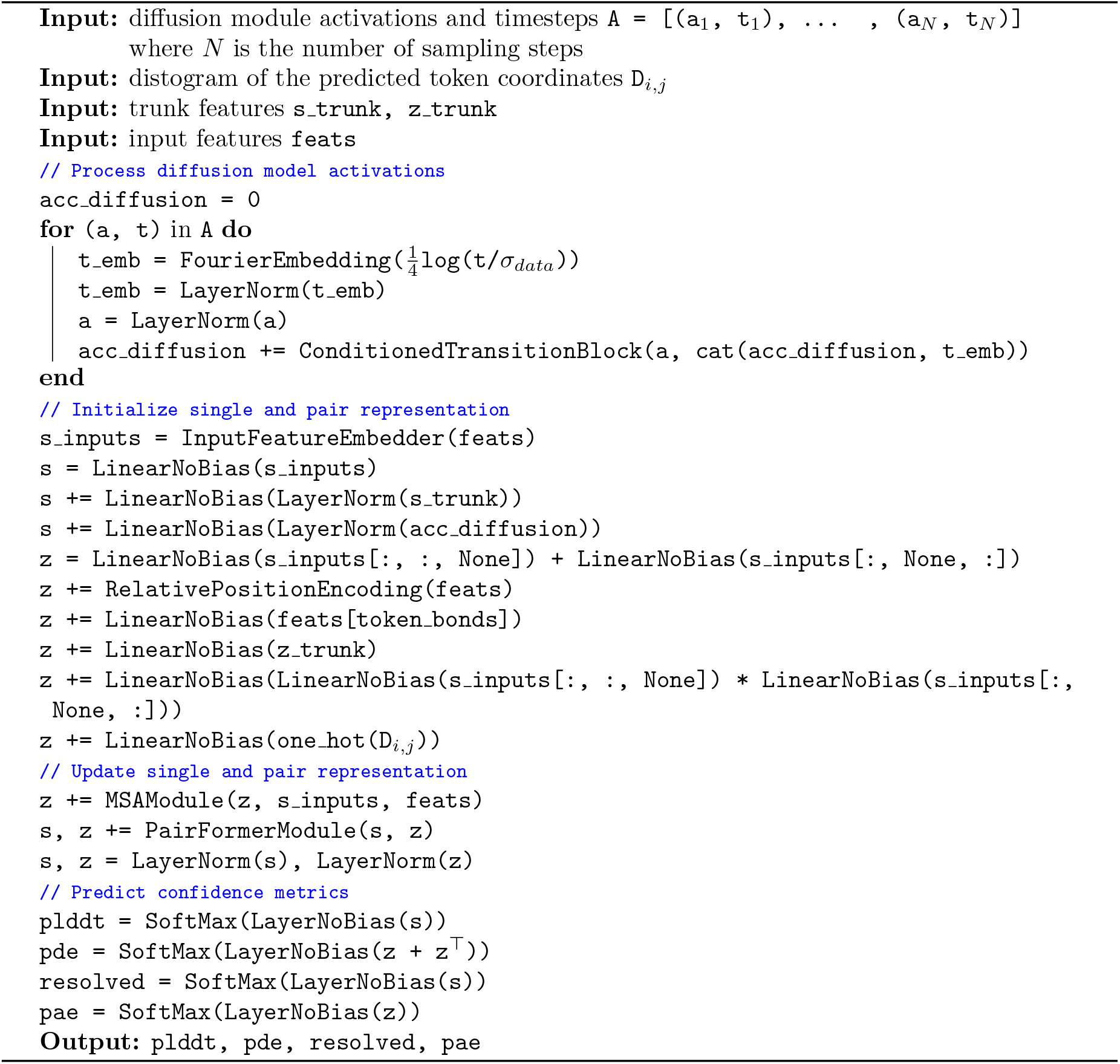

### 3.4 Optimizations

Below we summarize some computational techniques we use to speed up and/or reduce the memory consumption of the model. For details on the implementation of each of these, please refer to our code repository.

#### Sequence-local atom representation

The `AtomAttentionEncoder` and `AtomAttentionDecoder` include a pair-biased transformer on the representations of every atom. In particular, the attention of these transformers is sequence-local: blocks of 32 atoms only attend to the 128 atoms that are closest to them in sequence space. We developed a GPU-efficient implementation of the sequence-local attention precomputing a mapping (performed in blocks of 16 tokens) to the key and query sequence embeddings for each 32 key tokens block. The attention is then performed in parallel in each 32×128 block achieving a block sparse attention with dense matrices.

#### Attention bias sharing and caching

At a high level the denoising model must be run repeatedly for every diffusion timestep for every separate sample we take, while the trunk can be run once and its representation fed to all those denoising model passes.

The most expensive components of the denoising model are represented by the computation of the attention pair bias for the token and atom transformers. However, by examining their computational graph, we find that these elements do not depend either on the particular input structure given to the denoising model or the diffusion timestep. In particular, these elements are: the attention bias of all the transformer layers in the `AtomAttentionEncoder`, `AtomAttentionDecoder`, and `DiffusionTransformer`, and the intermediate single and pairwise atom representations of the `AtomAttentionEncoder`. Therefore, we can also run these components once and share them across all the samples and the entirety of the reverse diffusion trajectory, significantly reducing the computational cost of the reverse diffusion at the cost of storing these representations and biases in memory.

#### Greedy symmetry correction

During validation and confidence model training, the optimal alignment between the ground truth and predicted structure must be determined, accounting for permutations in the order of identical chains or symmetric atoms within those chains. Because the number of possible perturbations grows exponentially with the size of the complex, considering all of them is computationally unfeasible.

We devise the following procedure to perform an approximate, yet effective, atom matching. This operates in a hierarchical way searching (1) the optimal assignment of chains, and, then, (2) assuming chain assignment, to select atom perturbations greedily for every ligand or residue. For the first, for each symmetric chain, we compute the resulting global LDDT without changing any inner atom assignment. For the second step, iteratively for every ligand, amino acid, or nucleotide basis (one at a time), we find the perturbation of that ligand the most improves the global LDDT and greedily apply it.

Note that because the LDDT between pairs of elements that were not affected by the perturbation does not change, one can implement the test in the last step very efficiently only looking at the specific rows and columns of the distance matrix that change. In practice, we limit the number of perturbations of the chain assignment we consider to 100 and the perturbations of the atoms of each ligand to 1000.

#### Chunking for the MSA Module and Triangular Attention

To optimize memory efficiency in our model, we implement a chunking strategy that significantly reduces peak memory usage during inference. We follow the OpenFold chunking implementation for the Triangular Attention layer [Ahdritz et al., 2024], and extend it to the MSA Module, applying it at three critical points: the transition layers, pair-weighted average layers, and outer product layers. This improvement ensures the scalability of our model to larger inputs while maintaining a similar speed.

#### Trifast kernel

Triangle Self Attention, used in the PairFormer, suffers from *O*(*n*^3^) memory and time complexity. The memory complexity is often the limiting factor with regards to input size, for a given GPU. At moderate input sizes, it also dominates the runtime. A common solution to the memory issue is applying chunking to the calculation, which is typically done in PyTorch. We implement a Triton kernel that performs the calculation via a chunked online softmax. As such, the memory complexity is improved to *O*(*n*^2^). It also exploits the GPU architecture to speed up the computation for practical input sizes. This is an extension of FlashAttention that allows for the inner bias term [Dao et al., 2022]. A plot of the forwards runtime, compared to compiled PyTorch and another efficient kernel implementation [Song et al., 2023] can be found in Figure 4. The implementation can be found at https://github.com/latkins/trifast.

**Figure 4:**
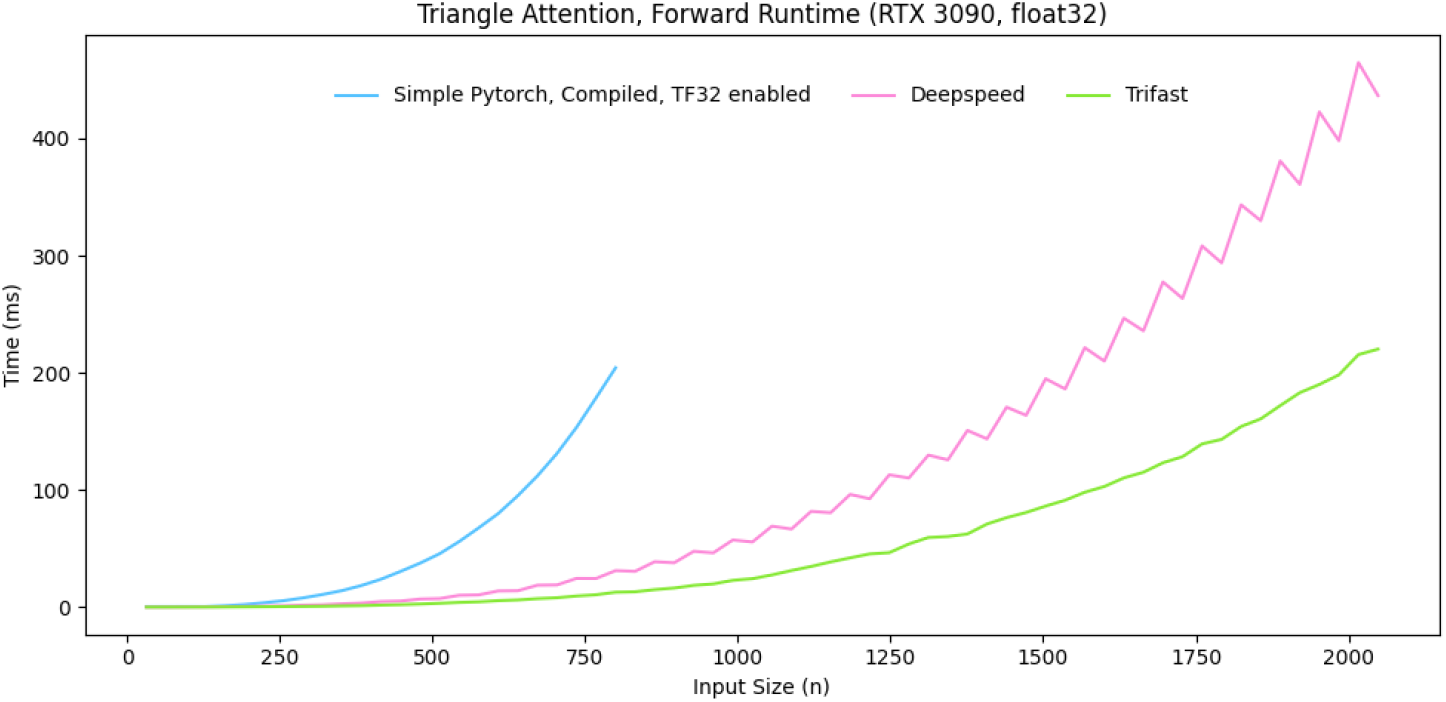
Forwards runtime of trifast kernel compared to compiled PyTorch and DeepSpeed kernel.

## 4 Boltz steering

### 4.1 Introduction

A visual inspection of several predictions from Boltz-1 revealed instances of hallucinations in the model”s outputs. The most prominent type of hallucination involved the placement of entire chains directly on top of one another. Abramson et al. [2024] noted a similar behaviour with AlphaFold3. Moreover, like previous machine learning-based structure prediction methods [Buttenschoen et al., 2024], we observe some non-physical structures predicted by the model. Examples of these behaviours include steric clashes between atoms, slightly incorrect bond lengths and angles, incorrect stereochemistry at chiral centers and stereobonds and aromatic rings predicted to be non-planar. While these issues are not strongly penalized in geometric measures of accuracy of the poses, they can prevent the predictions from being used in many downstream applications, both for expert and computational analyses (e.g. running molecular dynamics calculations). To tackle these issues, we introduce a new inference time steering technique that we refer to as Boltz-steering.

We aim to fix these issues by tilting the distribution defined by the trained diffusion model with a newly defined physics-inspired potential function.

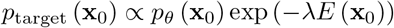

where *p*_*θ*_ is the data distribution and *E* the potential.

To sample from this tilted distribution, one approach would be to sample k particles from our diffusion process 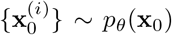, and subsequently resample them based on their corresponding importance weights exp 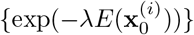. This approach would function similarly to the approach employed by Abramson et al. [2024] in AlphaFold3 where multiple samples of the diffusion model are then filtered based on physical-realism heuristics. However, as noted by Singhal et al. [2025], this approach has drawbacks. First, for constraints that are rarely satisfied by the model, importance sampling or filtering will be unable to bias the model towards the rare conformer that satisfies the constraint.

#### Related work

In the space of biomolecular structures, there have been other works using inference time potentials to guide the diffusion process. RFDiffusion [Watson et al., 2023] computes potentials based on the **x**_**0**_ prediction at each time step and applies a gradient update to encourage specific symmetric protein motifs and inter-chain contacts. Ishitani and Moriwaki [2025] also applies gradient updates to improve ligand structures using an RDKit conformer to define loss functions based on the RMSE between the bond lengths, bond angles, and chiral volumes of the predicted **x**_**0**_ conformer and reference conformer. By contrast, we use a flat-bottom potential based on the distance bounds, to enforce that the potential is within the range of realistic conformers without forcing it to match the RDKit conformer, expand to other physical properties and not only use gradient updates but also resampling. Finally, DecompDiff [Guan et al., 2024] also uses gradient updates in the reverse diffusion to improve the physical quality of generated molecule conformers.

### 4.2 Method

Boltz-steering takes a different and more effective approach, built on top the Feynmac-Kac (FK) steering framework introduced by Singhal et al. [2025]. We employ potential functions *G*_*t*_ (**x**_*T*_, …, **x**_*t*_) to tilt the transition kernels of the diffusion process *p*_*θ*_ (**x**_*t−*1_ | **x**_*t*_) at each intermediate timestep such that the trajectory is biased towards paths where the eventual **x**_0_ will have a low energy - or, equivalently, as described in the initial formulation, a large reward value.

During the steered reverse diffusion process, we begin with a sample from the initial tilted distribution at time *t* = *T*

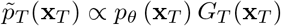

and at each subsequent timestep *t < T*, we sample from the tilted transition kernel

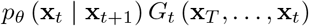

We define *G*_*t*_ in terms of the energy difference between the predicted denoised conformer 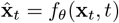 at subsequent timesteps.

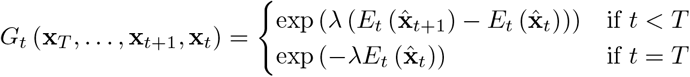

*E*_*t*_ is a weighted sum of constraint potentials, each tackling a specific physical issue. For each of these constraint potentials *con*, is defined as a flat-bottomed energy function *E*_*con*_(**x**), such that the energy *E*_*con*_(**x**) = 0 for any conformer **x** that satisfies the constraint and *E*_*con*_(**x**) is increasingly positive for conformers that violate the constraint to a greater degree. These potentials are summed to form *E*_*t*_ with time-dependent weights as the different physical errors of the model occur at different timescales (e.g. overlapping chains are set at high noise levels, strained bonds at low noise levels).

Directly sampling from the tilted transition kernel is intractable, therefore, the FK steering framework uses Sequential Monte Carlo (SMC) to propose multiple particles at each timeste,p which are then resampled based on their importance weights. For ease of notation, we use 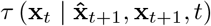 to denote the sampling algorithm used by Boltz at timestep t, such that the transition distribution of the diffusion model corresponds to

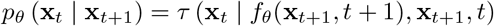

We further bias the diffusion trajectory towards low-energy conformers by incorporating guidance. Specifically, we employ backwards universal guidance introduced by Bansal et al. [2023], and define our proposal distribution as

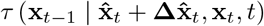

where 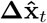 is computed by taking m steps of gradient descent with respect to 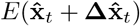 To adapt to these gradient steps we adjust the importance weights to be:

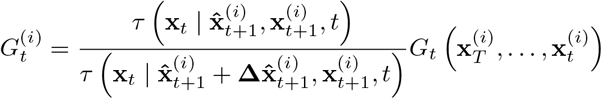

However, in practice we incorporate guidance at each time step while only resampling every 3 timesteps to facilitate sufficient exploration. The full Boltz-steering algorithm is presented in Algorithm 2.

### 4.3 Constraint potentials

We define the overall potential *E*_*t*_ as a linear combination of the various constraint potentials.

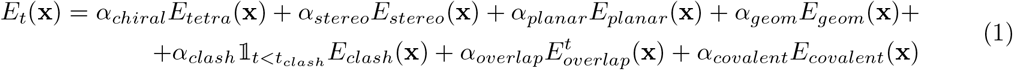

The clash potential is only applied for *t < t*_*clash*_. Below we present the formalization of each of the constraint potentials.

#### Algorithm 2: Boltz-steering

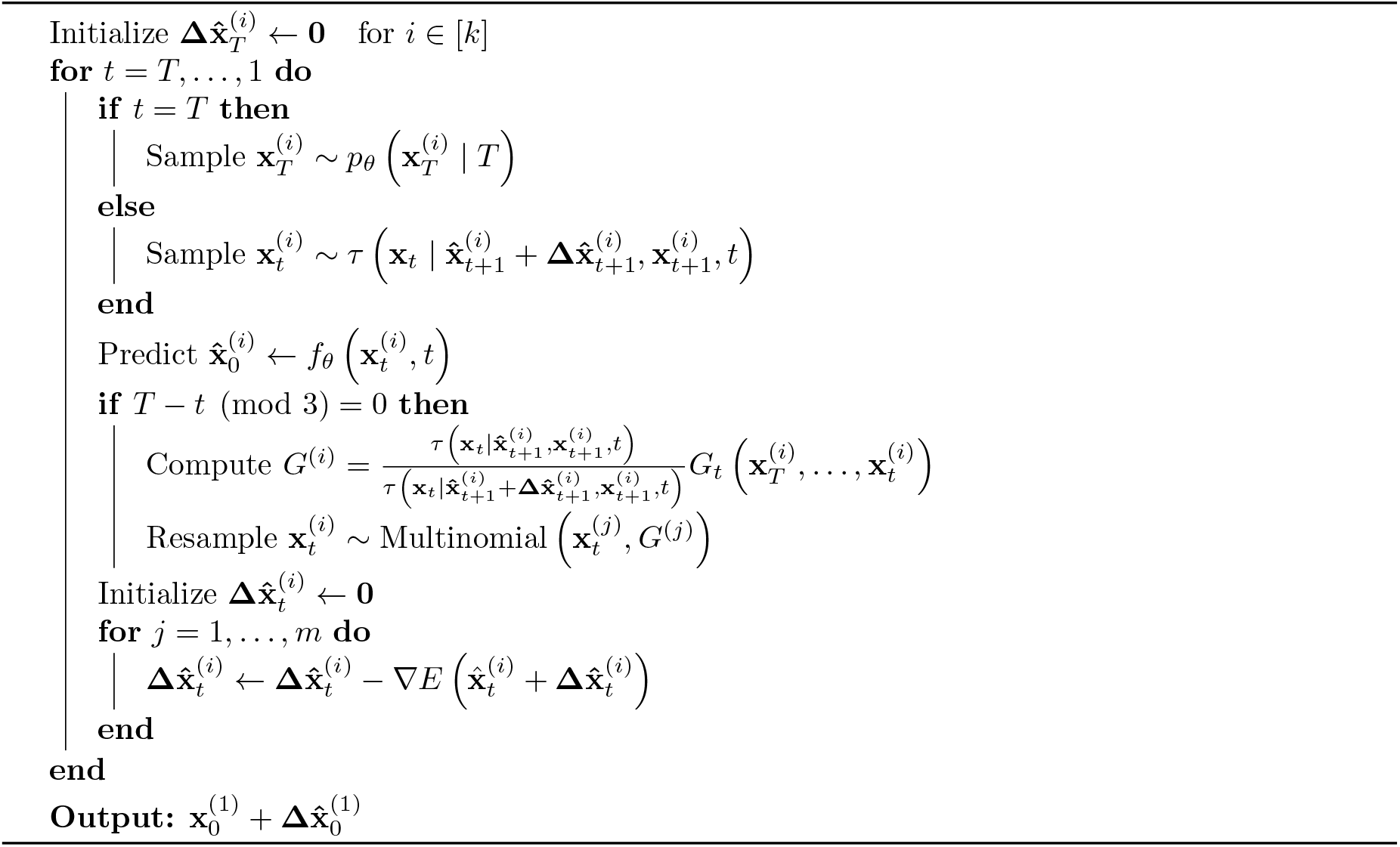

#### Tetrahedral Atom Chirality

For a chiral center Z with substituents A, B, C, D in decreasing Cahn-Ingold-Prelog (CIP) priority order, we say that Z has R chirality if the bonds (Z-A, Z-B, Z-C) form a right handed system and S chirality if they form a left handed system. To enforce that the predicted conformers have the correct chirality, we define potentials based on the improper torsion angles (X1, X2, X3, Z).

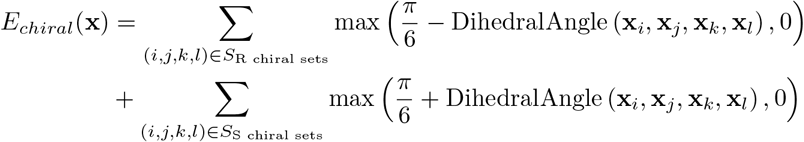

#### Bond Stereochemistry

For a bond Z1=Z2 where Z1 has substituents A1, B1 and Z2 has substituents A2, B2 in decreasing CIP priority order, we say that Z1=Z2 has E stereochemistry if the higher priority atoms A1 and A2 are on opposite sides of the bond and Z stereochemistry otherwise. To enforce that the predicted conformers have the correct stereochemistry, we define potentials based on the torsion angles (A1, Z1, Z2, A2) and (B1, Z1, Z2, B2).

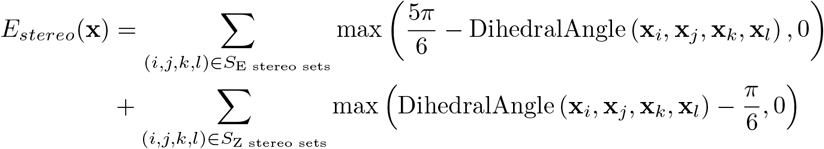

#### Planar Double Bonds

For planar double bonds C1=C2 between carbon atoms where C1 has substituents A1, B1, and C2 has substituents A2, B2, we define a flat bottom potential function to enforce planarity based on the improper torsion angles (A1, B1, C2, C1) and (A2, B2, C1, C2).

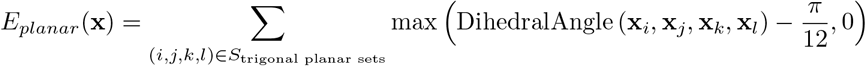

#### Internal Geometry

To ensure that the model generates ligand conformers with a physically realistic distance geometry, we define a flat-bottomed potential based on the bounds matrix which is generated by the RDKit package. For a ligand with *N* atoms, we define the lower and upper bounds matrices as *L, U ∈* ℝ^*N×N*^ where for a pair of atoms (i, j), the lower and upper distance bounds are given by *L*_*i,j*_, and *U*_*i,j*_ respectively [Buttenschoen et al., 2024]:

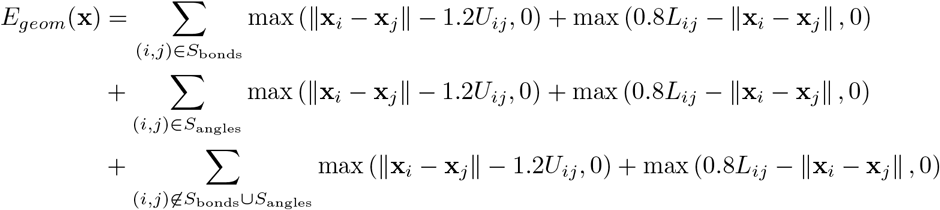

#### Steric Clash

To prevent steric clashes, we constrain the distance between atoms in distinct and non-bonded chains to be greater than 0.725 times the sum of the atoms’ Van der Waals radii.

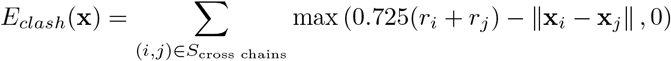

where the Van der Waals radius of atom i is denoted by *r*_*i*_

#### Overlapping Chains

To prevent overlapping chains, we define a time dependent potential based on the distance between the centroids of symmetric chains with more than one atom.

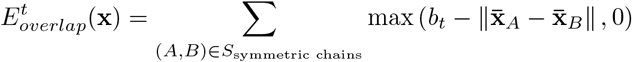

where 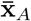 is the centroid of chain A and *b*_*t*_ is a time dependent parameter which controls the minimum distance between centroids and smoothly interpolates between 5.0 Å at *t* = 1 and 1.0 Å at *t* = 0 according to the schedule

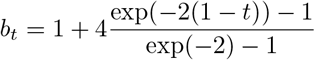

#### Covalently Bonded Chains

To ensure that the model respects covalently bonded chains, we define a potential to enforce that covalently bonded atoms between separate chains are within 2Å.

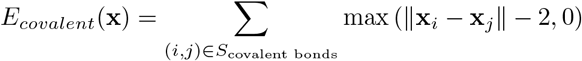

## 5 Results

We evaluate the performance of the model on two benchmarks: the diverse test set of recent PDB structures that we cured as discussed in Section 2.2, and CASP15, the last community-wide protein structure prediction competition where for the first time RNA and ligand structures were also evaluated [Das et al., 2023, Robin et al., 2023]. Both these benchmarks contain a very diverse set of structures including protein complexes, nucleic acids, and small-molecule, making them great testbeds for the assessment of models, such as Boltz-1, capable of predicting the structure of arbitrary biomolecules.

### Benchmarks

For CASP15, we extract all the competition targets with the following filters: (1) they were not canceled from the competition, (2) they have an associated PDB id to obtain the ground truth crystal structure, (3) the number of chains in the stochiometry information matches the number of provided chains, (4) the total number of residues with below 2000. This leaves a total of 76 structures. For our test set, we remove structures with covalently bounded ligands because the current version of the Chai-1 public repository does not provide a way to set these. Finally, for both datasets, we remove structures that go out of memory or fail for other reasons for any of the methods on A100 80GB GPUs. After these steps, we are left to evaluate 66 structures for CASP15 and 541 structures for the test set.

### Baselines

We evaluate our performance against AlphaFold3 [Abramson et al., 2024] and Chai-1 [Chai et al., 2024], current state-of-the-art biomolecular structure prediction models that were released under an exclusive commercial license and do not have training code and pipelines available. We ran the Chai-1 model using the `chai_lab` package version 0.2.1.

All the models were run with 200 sampling steps and 10 recycling rounds, and producing 5 outputs. We also used the same pre-computed MSA”s up to 16384 sequences. Since Chai-1 requires annotating the source of the sequences, we annotated all Uniref sequences with the `uniref90` label and all other sequences with the `bfd_uniclust` label. We briefly experimented with alternative labelings but did not find these to impact the model substantially.

### Evaluation criteria

We consider several well-established metrics to evaluate the performance of the models on these very diverse sets of biomolecules and structures. In particular, we compute:

1. The mean all-atom LDDT: measuring accuracy of local structures across all biomolecules;
2. The average DockQ success rates, i.e. the proportion of predictions with DockQ *>* 0.23, which measures the number of good protein-protein interactions predicted;
3. The average protein-ligand interface LDDT: measuring the quality of the ligand and pocket predicted interactions, official CASP15 metric to evaluate the ligand category;
4. The proportion of ligands with a pocket-aligned RMSD below 2Å: a widely adopted measure of molecular docking accuracy.
5. The physical quality of the poses generated by the different models by looking at the proportion of poses that pass a set of physical rules from PoseBusters [Buttenschoen et al., 2024]. In particular, we look at:
  a. Whether ligand bonds are strained by checking if the distances between bonded atoms are within a realistic range determined by RDKit
  b. Whether ligand angles are strained by checking if the 1-3 distances between atoms are within a realistic range determined by RDKit
  c. Whether ligand internal clashes are present by checking if the distances between all other pairs of atoms are above a lower bound determined by RDKit
  d. Whether tetrahedral atom chirality is preserved
  e. Whether bond stereochemistry is preserved
  f. Whether inter-chain clashes exist by checking if the distances between atoms in distinct and non-bonded chains with more than one atom are greater than 0.75 times the sum of their Van der Waals radii.

All metrics were computed using OpenStructure [Biasini et al., 2013] version 2.8.0. LDDT-PLI, DockQ and ligand RMSD success rates are computed over all the different protein-protein and protein-ligand interfaces, these proportions are averaged over interfaces within individual complexes and then averaged across complexes containing interfaces. Following a similar format to that used in CASP and to allow a fair comparison of the methods, we run all methods to generate 5 samples and evaluate both the best (oracle) and highest confidence prediction (top-1) out of the 5 for every metric.

To foster further development of methods and the convergence of the field towards well-curated and adopted benchmarks, we publicly release all the inputs, outputs, and evaluations of all the models in our benchmarks as well as the scripts we used to aggregate them. The instructions for downloading them are available on our GitHub repository^4^.

### Results

We report the performance of AlphaFold3, Chai-1 and Boltz-1 in Figures 5 and 7. Overall the models show comparable results across the different metrics across both CASP15 and the test set.

**Figure 5:**
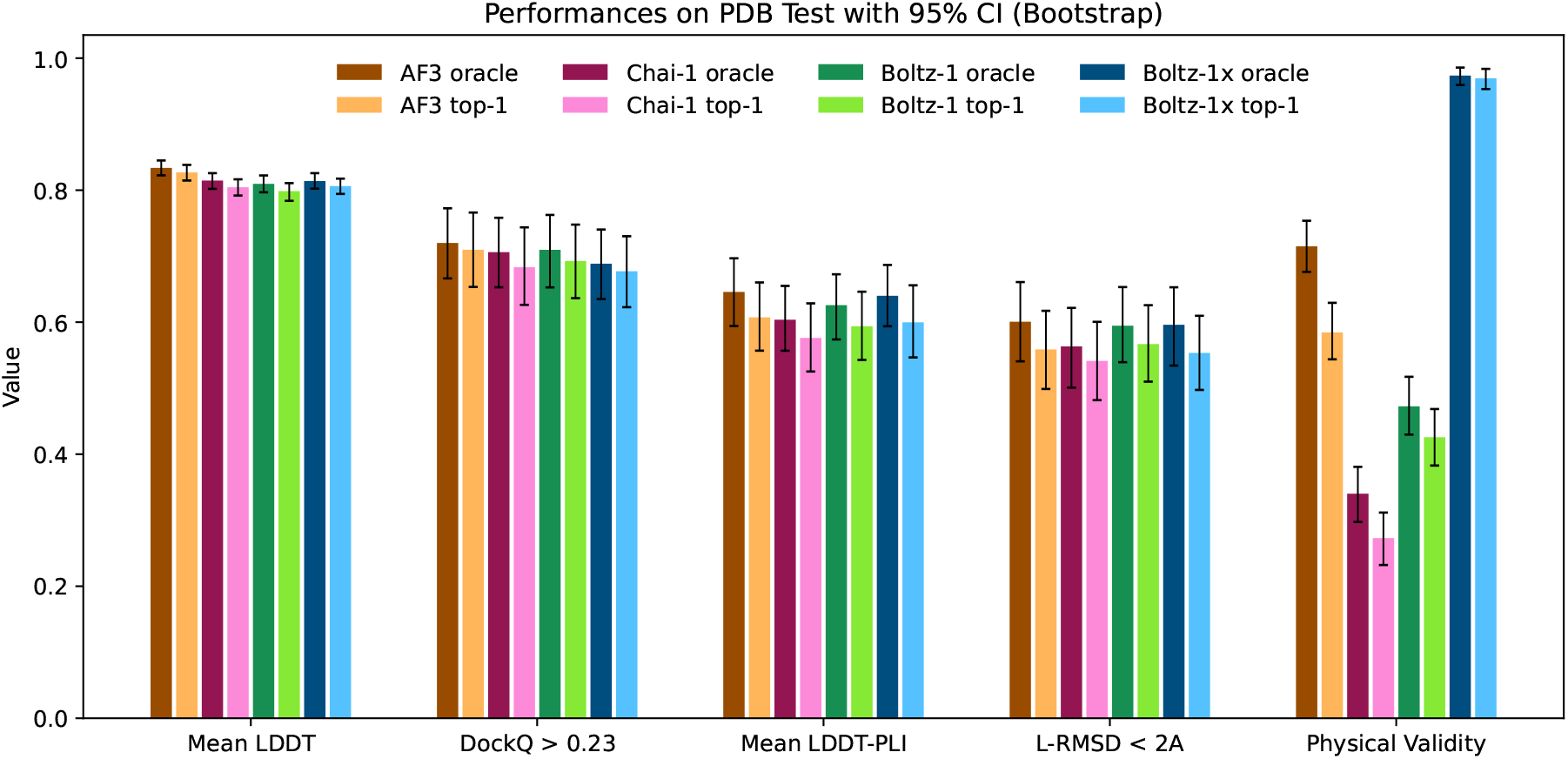
Visual summary of the performance of AlphaFold3, Chai-1, Boltz-1 and Boltz-1x on the test set.

**Figure 6:**
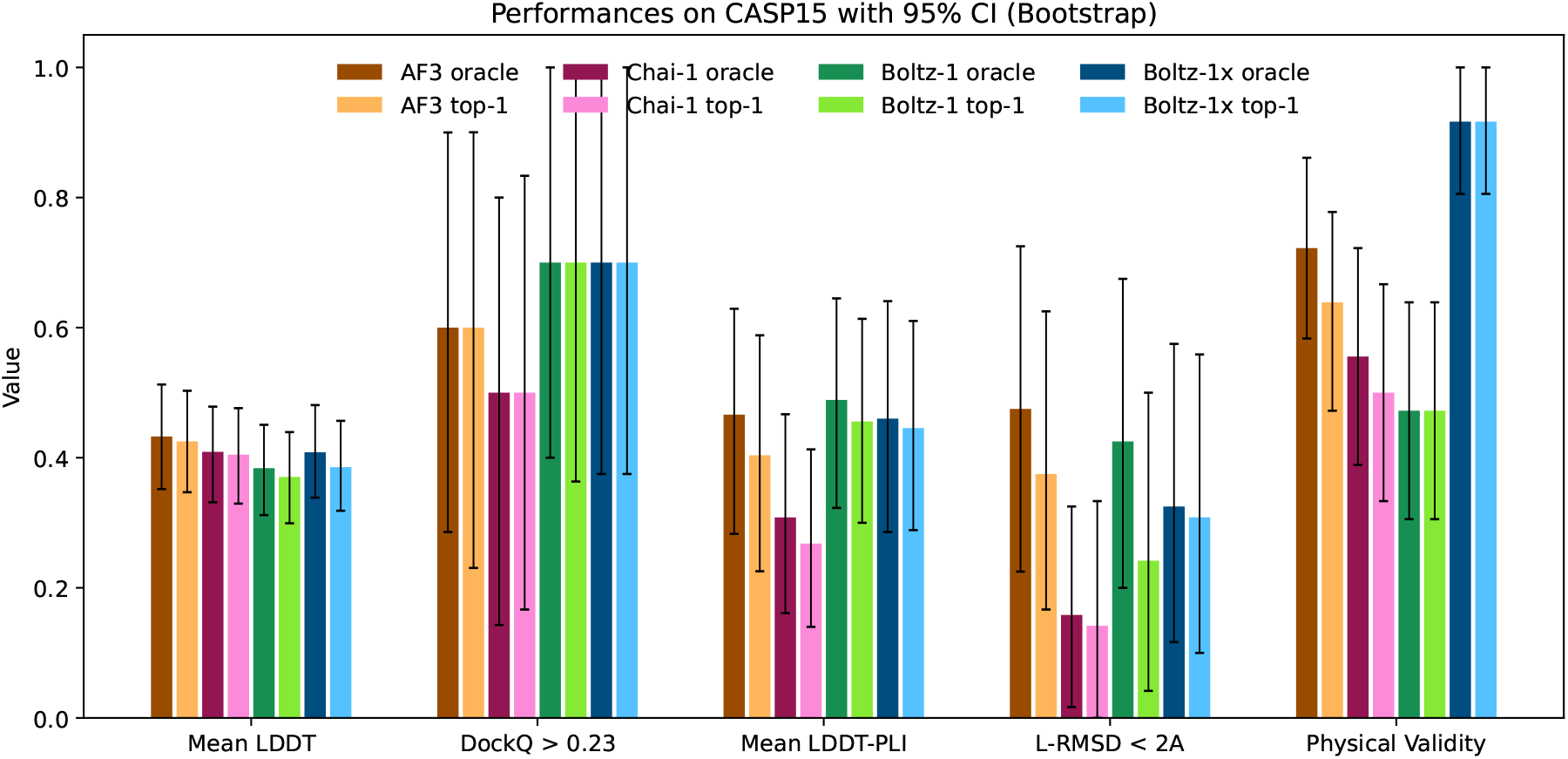
Visual summary of the performance of AlphaFold3, Chai-1, Boltz-1 and Boltz-1x on the CASP15 benchmark.

**Figure 7:**
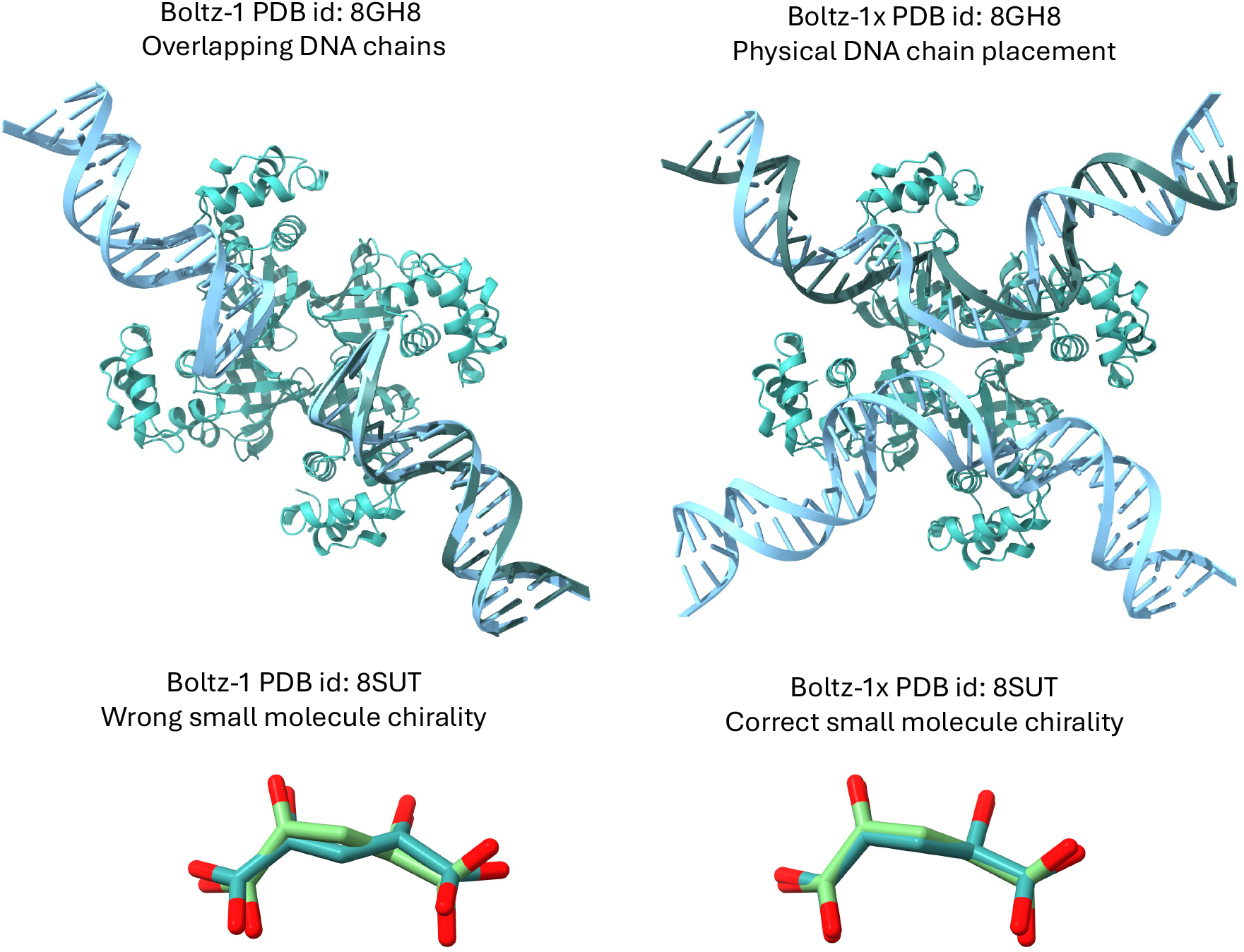
Examples of some failure modes of Boltz-1 leading to unphysical poses, on the left, and the fixed poses resulting from Boltz-1x, on the right.

AlphaFold3 has a slight edge over the other models on the mean LDDT metric, which likely derives from better handling complexes containing RNA and DNA thanks to its extra distillation datasets.

For protein-protein interactions, the performance of the methods is also aligned. AlphaFold3 slightly outperforms Boltz-1 and Chai-1 on the test set in terms of proportion of interfaces with DockQ *>* 0.23, however, all differences are well within the confidence intervals. Similarly, in the protein-ligand metrics, AlphaFold3 and Boltz-1 obtain slightly better mean LDDT-PLI and proportion of ligand RMSD < 2Å than Chai-1, but once again these differences are within the confidence intervals. These results demonstrate, especially in terms of the accuracy of predictions for protein-protein and protein-ligand interactions that Boltz-1 obtains a performance comparable to that of the state-of-the-art models AlphaFold3 and Chai-1.

When evaluating Boltz-1 on the physical quality tests we see that 57% of top-1 poses do not pass these tests on the test set, suggesting that they likely have severe physical issues. Chai-1 and AlphaFold3 also have relatively low rates of complexes passing all the checks, with respectively around 27% and 58%. On the other hand, Boltz-1x gets 97% of the poses passing the checks, while maintaining a similar level of performance compared to the other models.

In Table 1, we report a number of ablations with respect to the number of recycling steps and diffusion steps. These show a generally monotonic improvement in the performance with more steps, which is relatively plateaued beyond 3 recycling and 50 diffusion steps. Finally, in Figure 1 we present two examples of hard targets from the test set where Boltz-1 performed remarkably well with TM scores around 95%

**Table 1:** Ablation on the number of recycling rounds and sampling steps for Boltz-1 on the test set. We run the ablation study generating 5 samples and evaluating both the best (oracle) and highest confidence prediction (top-1) out of the 5 for every metric. All models used pre-computed MSAs with up to 4,096 sequences. It is worth noting that the metrics are noisy, so minor inconsistencies (e.g., lack of improvement with increased recycling rounds or diffusion steps) should not be overinterpreted. Moreover, there is a slight difference with the results in Figures 5 and 7 due to differences in MSA parameters as well as the set of structures passing all ablations.

| # rec. | # steps | Mean LDDT |  | DockQ > 0.23 |  | Mean LDDT-PLI |  | L-RMSD < 2Å |  |
| --- | --- | --- | --- | --- | --- | --- | --- | --- | --- |
|  |  | oracle | top-1 | oracle | top-1 | oracle | top-1 | oracle | top-1 |
| 3 | 200 | 0.729 | 0.716 | 0.654 | 0.625 | 0.621 | 0.580 | 0.581 | 0.545 |
| 0 | 200 | 0.698 | 0.681 | 0.579 | 0.544 | 0.573 | 0.530 | 0.582 | 0.541 |
| 1 | 200 | 0.718 | 0.702 | 0.656 | 0.635 | 0.623 | 0.573 | 0.588 | 0.535 |
| 2 | 200 | 0.726 | 0.710 | 0.651 | 0.632 | 0.616 | 0.581 | 0.587 | 0.546 |
| 3 | 200 | 0.729 | 0.716 | 0.654 | 0.625 | 0.621 | 0.580 | 0.581 | 0.545 |
| 6 | 200 | 0.732 | 0.714 | 0.644 | 0.635 | 0.630 | 0.593 | 0.595 | 0.555 |
| 8 | 200 | 0.733 | 0.717 | 0.644 | 0.633 | 0.630 | 0.584 | 0.588 | 0.545 |
| 10 | 200 | 0.735 | 0.720 | 0.644 | 0.631 | 0.619 | 0.577 | 0.575 | 0.541 |
| 3 | 20 | 0.720 | 0.693 | 0.615 | 0.592 | 0.577 | 0.547 | 0.550 | 0.532 |
| 3 | 50 | 0.727 | 0.710 | 0.645 | 0.627 | 0.621 | 0.579 | 0.586 | 0.540 |
| 3 | 200 | 0.729 | 0.716 | 0.654 | 0.625 | 0.621 | 0.580 | 0.581 | 0.545 |

## 6 Conclusion

We introduced Boltz-1, the first fully commercially accessible open-source model to achieve AlphaFold3-level accuracy in predicting the 3D structures of biomolecular complexes. To accomplish this, we replicated and expanded upon the AlphaFold3 technical report, incorporating several innovations in architecture, data curation, training, and inference processes. We empirically validated Boltz-1 against AlphaFold3 and Chai-1, the current state-of-the-art structure prediction methods, demonstrating comparable performance on both a diverse test set and the CASP15 benchmark.

Further, we introduced Boltz-1x an updated model that leverages Boltz-steering, a new inference time technique, to significantly improve the physical quality of the poses generated while maintaining their geometric accuracy of Boltz-1.

The open-source releases of Boltz-1 and Boltz-1x represent significant steps forward in democratizing access to advanced biomolecular modeling tools and improving their applicability across domains. By freely providing the training and inference code, model weights, and datasets under the MIT license, we aim to enable researchers and organizations to experiment and innovate using Boltz-1 and Boltz-1x. We envision Boltz models as a foundational platform for researchers to build upon, fostering collaboration to advance our collective understanding of biomolecular interactions and accelerating breakthroughs in drug design, structural biology, and beyond.

## 7 Acknowledgments

We would like to thank Sergey Ovchinnikov, Bowen Jing, Hannes Stark, Jason Yim, Peter Mikhael, Richard Qi, Wengong Jin, Rohith Krishna, Evan Feinberg, and Maruan Al-Shedivat for the invaluable discussions and help. We also thank the research community for all the feedback we received, that has helped us improve the usability of the model, understand its limitations, and help inform improvements that we are doing for future versions of the model.

Large portions of the GPU resources necessary to complete the project were provided by Genesis Therapeutics and the US Department of Energy. For the latter, we acknowledge our use of the National Energy Research Scientific Computing Center (NERSC), a Department of Energy Office of Science User Facility, via NERSC award GenAI@NERSC. This work was also supported by the NSF Expeditions grant (award 1918839: Collaborative Research: Understanding the World Through Code), the Abdul Latif Jameel Clinic for Machine Learning in Health, the DTRA Discovery of Medical Countermeasures Against New and Emerging (DOMANE) Threats program, and the MATCHMAKERS project supported by the Cancer Grand Challenges partnership financed by CRUK (CGCATF-2023/100001) and the National Cancer Institute (OT2CA297463).

## Algorithm 3: Dense MSA Pairing

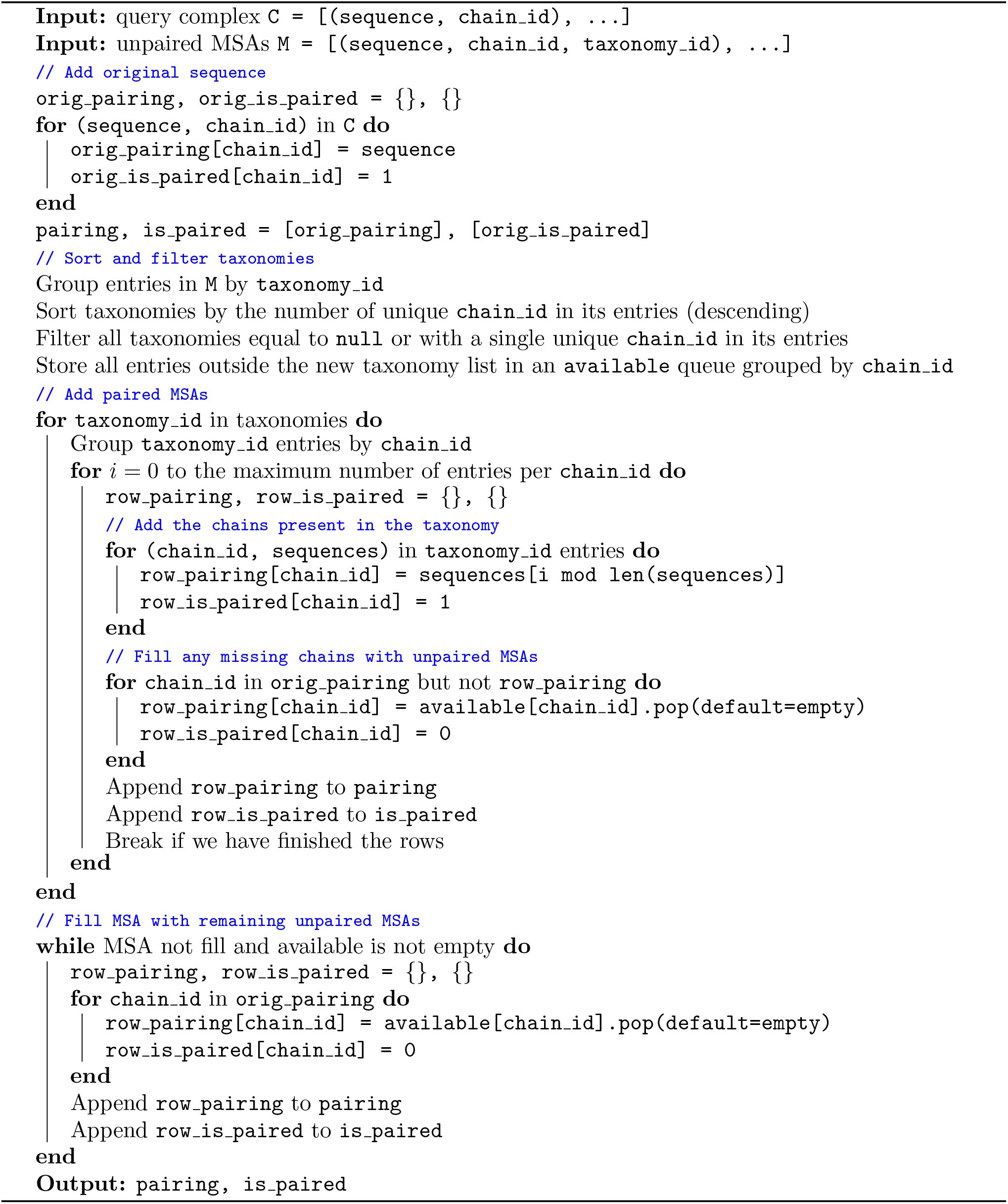

## Algorithm 4: Unified Cropping

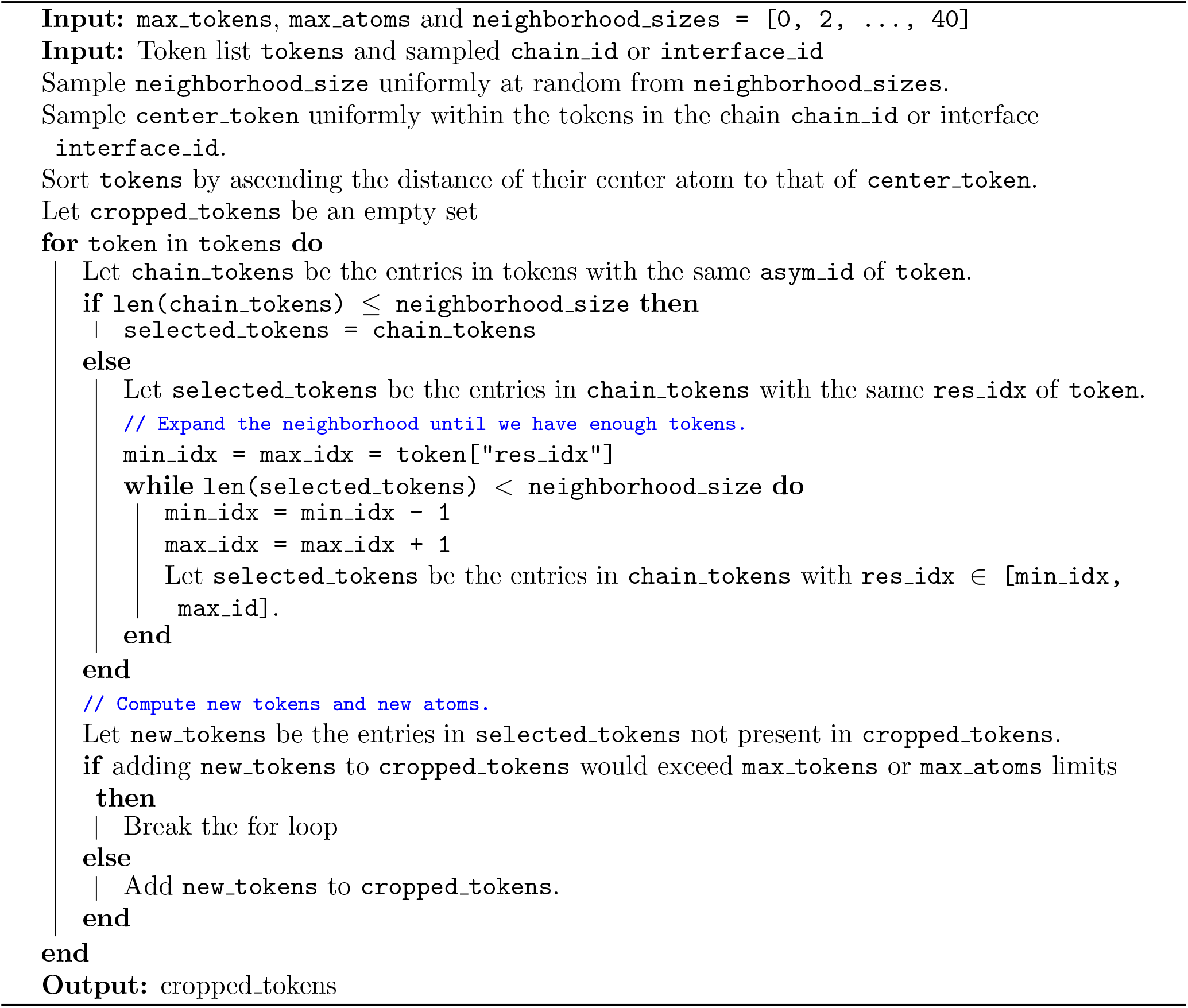

## Algorithm 5: Robust pocket-conditioning

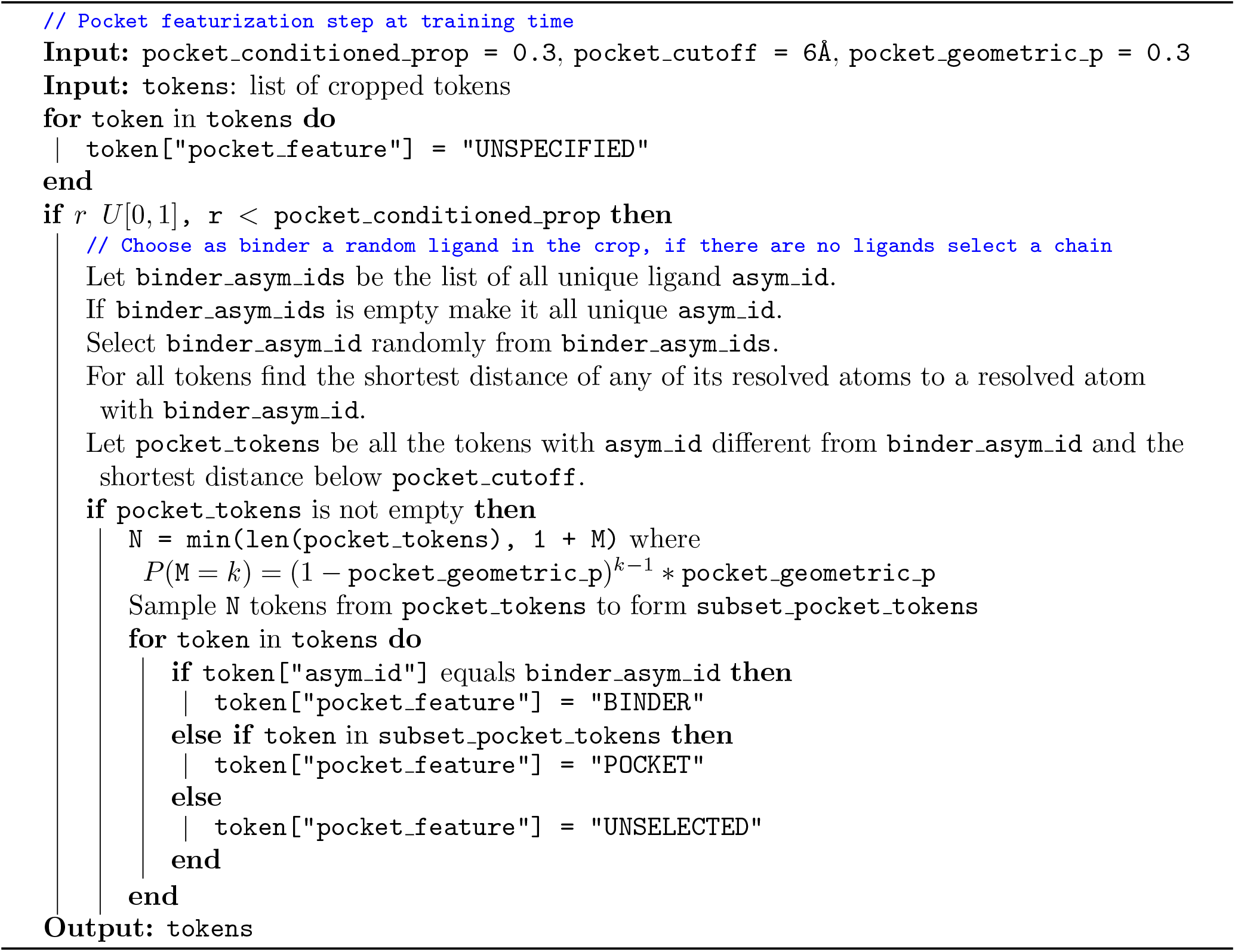

## Footnotes

1 https://github.com/jwohlwend/boltz

2 Some of these differences may simply be the result of reporting mistakes in the current version of the original manuscript from Abramson et al. [2024], as reported.

3 We note that a similar strategy was also concurrently noticed by https://github.com/Ligo-Biosciences/AlphaFold3.

4 https://github.com/jwohlwend/boltz

